# Vessel Density Mapping of Cerebral Small Vessels on 3D High Resolution Black Blood MRI

**DOI:** 10.1101/2023.03.18.533300

**Authors:** Mona Sharifi Sarabi, Samantha J. Ma, Kay Jann, John M. Ringman, Danny J.J. Wang, Yonggang Shi

**Affiliations:** USC Stevens Neuroimaging and Informatics Institute, University of Southern California, Los Angeles, CA, USA; Siemens Healthcare USA, Los Angeles, CA, USA; Department of Neurology, University of Southern California, Los Angeles, CA, USA

**Keywords:** High-resolution black-blood MRI, Turbo spin-echo with variable flip angles (TSE VFA), vessel segmentation, vessel density, small vessel disease

## Abstract

Cerebral small vessels are largely inaccessible to existing clinical in vivo imaging technologies. This study aims to present a novel analysis pipeline for vessel density mapping of cerebral small vessels from high-resolution 3D black-blood MRI at 3T. Twenty-eight subjects (10 under 35 years old, 18 over 60 years old) were imaged with the T1-weighted turbo spin-echo with variable flip angles (T1w TSE-VFA) sequence optimized for black-blood small vessel imaging with iso-0.5mm spatial resolution at 3T. Hessian-based vessel segmentation methods (Jerman, Frangi and Sato filter) were evaluated by vessel landmarks and manual annotation of lenticulostriate arteries (LSAs). Using optimized vessel segmentation, large vessel pruning and non-linear registration, a semiautomatic pipeline was proposed for quantification of small vessel density across brain regions and further for localized detection of small vessel changes across populations. Voxel-level statistics was performed to compare vessel density between two age groups. Additionally, local vessel density of aged subjects was correlated with their corresponding gross cognitive and executive function (EF) scores using Montreal Cognitive Assessment (MoCA) and EF composite scores compiled with Item Response Theory (IRT). Jerman filter showed better performance for vessel segmentation than Frangi and Sato filter which was employed in our pipeline. Cerebral small vessels on the order of a few hundred microns can be delineated using the proposed analysis pipeline on 3D black-blood MRI at 3T. The mean vessel density across brain regions was significantly higher in young subjects compared to aged subjects. In the aged subjects, localized vessel density was positively correlated with MoCA and IRT EF scores. The proposed pipeline is able to segment, quantify, and detect localized differences in vessel density of cerebral small vessels based on 3D high-resolution black-blood MRI. This framework may serve as a tool for localized detection of small vessel density changes in normal aging and cerebral small vessel disease.

## 1. Introduction

Cerebral small vessel disease (cSVD) is often associated with neurodegenerative diseases, and it can contribute to the progression of cognitive decline and physical disabilities (Breteler et al., 1994; Debette et al., 2010; Mayda and DeCarli, 2009; Rosenberg et al., 2016; Wardlaw et al., 2013a). Neuroimaging plays a key role in characterizing cSVD by identifying various features linked to cSVD including recent small subcortical infarcts, lacunes, white matter hyperintensities, enlarged perivascular spaces, microbleeds, and brain atrophy (Blair et al., 2017; Vemuri et al., 2022; Wardlaw et al., 2013b). However, when these features become visible or detectable in structural imaging, they are already manifestations of significant deterioration caused by cSVD. Recent advanced imaging techniques have facilitated the assessment of pathophysiology by determining microstructural tissue integrity (Baykara et al., 2016) as well as vascular function through the measurement of cerebral blood flow, arterial stiffness, cerebrovascular reactivity, and blood brain barrier leakage (Jann et al., 2021; Shao et al., 2019; Sur et al., 2020; Yan et al., 2016). Because the underlying mechanisms of cerebral microvascular changes in conditions related to cSVD remain poorly understood, there is an increasing desire to detect the early features of cSVD in order to prevent or mitigate downstream disease-related tissue changes.

Considering that most clinical features of vascular neurodegenrative diseases are consequences of cSVD, it is essential to be able to directly visualize and evaluate the cerebral small vessels themselves. To date, however, these small vessels including arterioles, capillaries and venules are largely inaccessible to existing clinical in vivo imaging technologies. We recently proposed and optimized high resolution 3D black-blood MRI with sub-millimeter (iso-0.5mm) spatial resolution using T1-weighted turbo spin-echo with variable flip angles (T1w-TSE-VFA) at field strengths of 3 and 7 Tesla (Ma et al., 2019). The long echo train of TSE-VFA offers two main advantages for visualizing small vessels: 1) adequate flow suppression by inherent dephasing of flowing signals (black-blood MRI); 2) sub-millimeter spatial resolution (isotropic ∼0.5–0.6 mm) and near whole-brain coverage in a clinically acceptable time (<10min). We demonstrated that optimized TSE-VFA is suitable for visualizing lenticulostriate arteries (LSAs) on the order of a few hundred microns, which can be manually segmented for the quantification of their branch number, length, and tortuosity (Cho et al., 2008; Kang et al., 2009; Ma et al., 2019).

The purpose of this study was to present a semiautomatic post-processing pipeline (**Figure 1**) for segmenting cerebral small vessels and further for mapping of small vessel density based on high resolution (isotropic ∼0.5mm) black-blood MRI with near whole-brain coverage at 3T. This pipeline included pre-processing steps (skull-stripping, bias correction and non-local means (NLM) denoising), followed by optimized vessel segmentation and non-linear registration for quantification of small vessel density across brain regions. The developed pipeline was evaluated by comparison of localized vessel density between young and aged groups, as well as association of vessel density in aged subjects with their neurocognitive scores at voxel-level.

**Figure 1.**
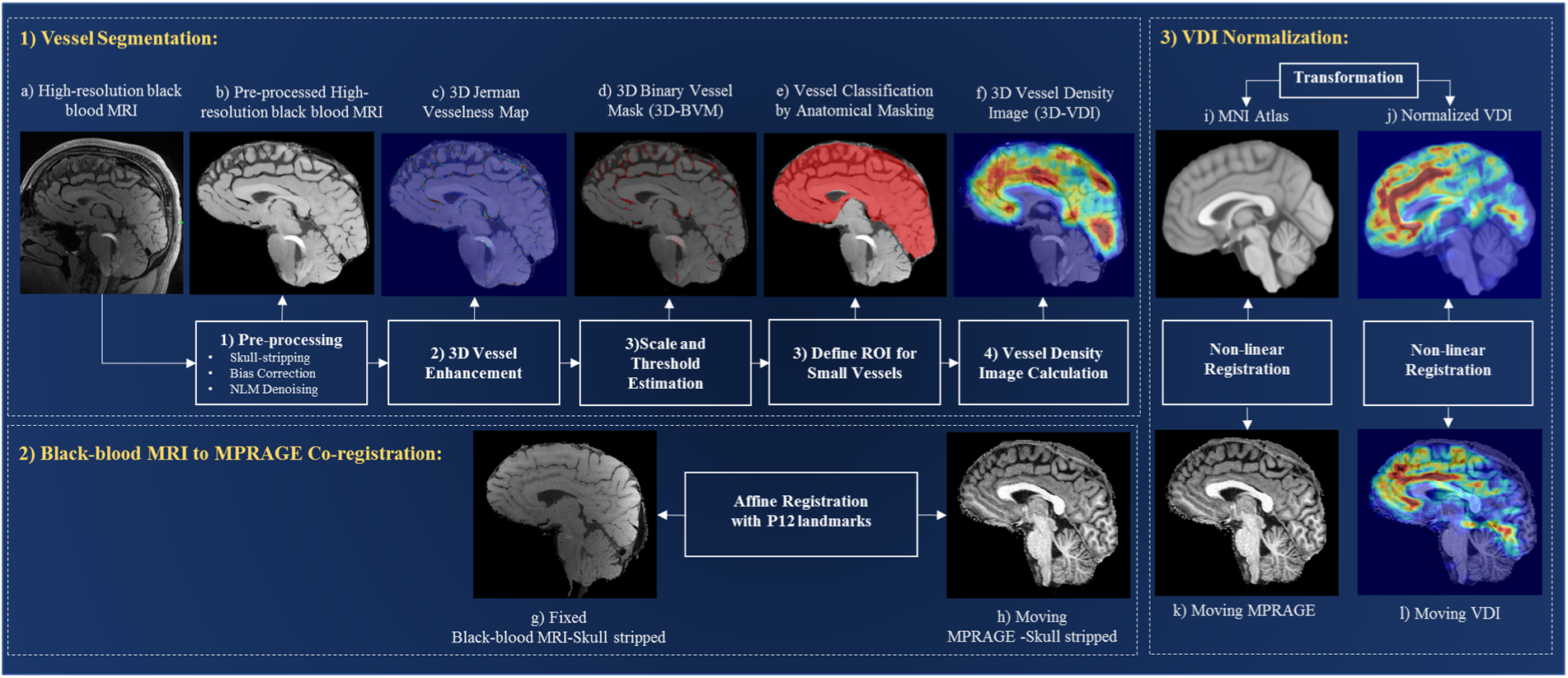
The proposed small vessel processing pipeline based on black-blood MRI. (1) Vessel segmentation: (a) Input high-resolution black-blood MRI to the pipeline. (b) Preprocessing steps such as skull-stripping, bias correction and denoising by non-local means (NLM) applied on (a). (c) 3D vessel enhancement, various multiscale Hessian-based methods (Frangi et al., 1998, Sato et al., 2000, Jerman et al., 2016) have been evaluated for enhancement of small vessels; Jerman filter outperformed other candidate methods. (d) Binarized vessel mask (BVM) was obtained by thresholding Jerman’s vesselness map using the optimal threshold. (e) Spatial masking was defined to construct region of interest (ROI) and classify small cerebral vessels. (f) Vessel density image (VDI) was calculated by diffusing the content of (e) to the entire image volume. (2) Co-registration of high-resolution black-blood MRI (g) to MPRAGE (h) for each subject with affine registration. 12 landmark points were incorporated to improve the robustness of co-registration. (3) VDI normalization: Mapping VDIs (l) from each subject to the MNI152 atlas (j) by combining the affine registration in (2) and the nonlinear registration between the MPRAGE (k) and the MNI152 atlas (i).

## 2. Methods

### 2.1 Subjects and Imaging

The study was approved by the institutional review board of the University of Southern California (USC) and in accordance with the principles of Declaration of Helsinki. Black-blood MRI and T1w structural images (MPRAGE) were collected from 28 participants including young control (N =10, 4 female, 27±3.5 years, age range [22,33]), and aged control (N =18, 14 female, 69.4±6 years, age range [60, 81]) groups. The subjects were screened for neurologic or psychiatric disorders and were generally healthy. In 12 of the 18 aged subjects, standard neuropsychological evaluation was performed. Images were acquired using a Siemens 3T MAGNETOM Prisma scanner with a 32-channel head coil. The “black blood” contrast was attained with an optimized T1w-TSE-VFA sequence (Ma et al., 2019) with the following parameters: TR/TE = 1000/12ms, turbo factor = 44, matrix size = 756×896, resolution = 0.51×0.51×0.64 mm^3^ interpolated to 0.5×0.5×0.5 mm^3^, 160 sagittal slices and 10% oversampling, GRAPPA factor = 2; total imaging time = 8:39 min. This sequence provided nearly whole brain coverage except that the sagittal FOV was 80mm, and bilateral saturation bands were applied to suppress the out-of-volume signals from temporal regions. Image quality grading of the high-resolution 3D black-blood MRI was performed by two expert graders. Grading criteria was based on a 4-point Likert scale, considering imaging artifacts such as motion, blurring, ringing, and wraparound. MRI images included in this study had an image quality grade and vascular quality grade of 3 or higher. This dataset will be referred to as high-resolution black blood-3T-28 (**HRBB-3T-28**) in the rest of the manuscript. Additionally, for method validation purposes, 15 black-blood images from the volunteer cohort of our previous LSA study (Ma et al., 2019) were used in this study. This cohort included a total of 15 healthy volunteers, with 9 participants (7 male, 27.2 ±3.5 years) between 22-33 years old and 6 participants (1 male, 64 ±2.4 years) older than 61 years of age, herein referred to as the young and aged group, respectively. All image volumes in this LSA dataset were separated into left and right hemispheres. This dataset will be referred to as **LSA-30** in the rest of the manuscript.

### 2.2 Pre-Processing

To prepare the images for vessel segmentation, initially, high-resolution black-blood MRI and MPRAGE images were skull-stripped using the Statistical Parametric Mapping (SPM) software. Due to intensity inhomogeneity and relatively low signal-to-noise ratio (SNR), skull-stripped images of high-resolution black-blood MRI were further pre-processed by bias correction and denoising via a non-local means (NLM) filter (Buades et al., 2005a). Due to the high computational cost of NLM in 3D (Buades et al., 2005b), block-wise NLM was adopted in this study which allows computationally efficient filtering without compromising the denoising result (Coupé et al., 2008).

In this study, the NLM search window size, which has an impact on computational time and visual quality of the results, was experimentally set to 3 × 3 × 3 in a block of size 32 × 32 × 32 within each 3D black-blood MRI volume. To avoid excessive denoising and preserve small vessel details, the smoothing parameter *h*, which depends on the noise level, was estimated as 0.03. **Figures S1(a)** and **(b)** in the supplementary material show an example of high-resolution black block MRI of a young control subject before and after pre-processing, respectively.

### 2.3 Vessel Enhancement in 3D

To overcome the undesired intensity variations of vessel images and suppress non-vascular structures, numerous filter-based vessel enhancement methods have been proposed (Agam et al. 2005, Wiemker et al. 2013, Krissian et al. 2000, Frangi et al. 1998, Sato et al. 2000, Li et al. 2003, Erdt et al. 2008, Zhou et al. 2007, Law and Chung 2010, Jiang et al. 2006, Zhang et al. 2010), and their performances have been significantly improved recently. Among these methods, Hessian-based filters which are based on second order image derivatives have shown great success in various clinical applications such as segmentation of liver vessels in abdominal CTA (Luu et al., 2015), lung vessels in thoracic CT (Rudyanto et al., 2014), and computer-aided detection of cerebral aneurysms in 3D rotational angiography, MRA and CTA (Hentschke et al., 2014). In this category of methods, one recent method called the *Jerman filter*, which is based on the ratio of Hessian matrix eigenvalues, has been proposed for enhancement of both 2D and 3D vasculatures (Jerman et al., 2016). The authors demonstrated that the Jerman filter could overcome some deficiencies of previous Hessian-based enhancement algorithms such as poor and non-uniform responses for vessels with varying contrast and sizes, bifurcations and aneurysms. The enhancement function in the Jerman filter is defined as:

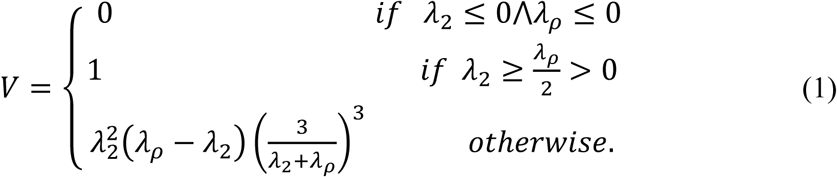

where, *λ*_2_ and *λ*_3_ are the eigenvalues of the Hessian matrix, and *λ*_*ρ*_ is the regularized value of *λ*_3_ and is defined below to ensure the robustness of vesselness measure to low magnitudes of *λ*_2_ *and λ*_3_:

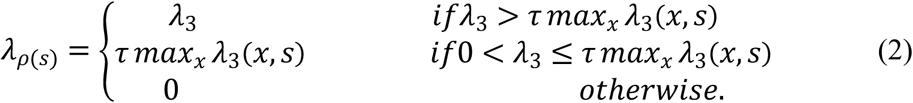

where, *τ* is a cutoff threshold between zero and one. Higher value of *τ* increases the difference between *λ*_2_ and *λ* magnitudes for low contrast structures. The proposed enhancement function’s response values range from 0 to 1.

In this study, according to the vascular anatomy of the brain and the study cohort that includes aged control subjects with thin, tortuous and low contrast vessels, an ideal enhancement function should exhibit robust, uniform, and high response in the presence of: i) varying noise and contrast; ii) varying vascular morphology (e.g., multiscale vessels especially small vasculature, vessels with different cross-sections (circular-elliptical)); iii) pathology (e.g., vessels thinning, high tortuosity or vessel bends). To meet these expectations, the Jerman filter appears to be a proper choice. To help justify our choice, the performance of the Jerman method will be compared with two widely used enhancement functions: the Frangi (Frangi et al., 1998) and Sato (Sato et al., 2000) method. We will present the comparison results among the candidate methods in **Section** 3.1.2.

Moreover, in multiscale Hessian-based methods, vessels’ scale range and the threshold value for binarizing the vesselness response are two important parameters to estimate. Inaccurate assessment of these parameters will result in over or under segmentation of the vascular network of interest. Therefore, to quantitatively evaluate the choice of parameters and compare the performance of the candidate methods, two sets of ground truth data were used. 1) manually delineated landmarks of cerebral small vessels from high-resolution black-blood MRIs (**Figure 2** (1)) that serve as the validation dataset for parameter optimization. This validation cohort was a subset of the HRBB-3T-28 subjects, comprised of five young controls and five aged control subjects. 2) 3D annotation of LSAs on unilateral LSA images of the LSA-30 dataset from the left and right hemispheres of 15 subjects (**Figure 2** (2)) were utilized as an independent testing dataset to further quantify the performance of the candidate vessel enhancement methods with optimized parameters estimated from the validation cohort. In the validation cohort, for each high-resolution black-blood MRI we annotated 2000 vascular and 2000 non-vascular landmarks (**Figure** 2) (drawn by experts with six years of experience in vascular image analysis). Vessel landmarks were mostly placed at the center and boundary of small and low contrast vasculature to increase the sensitivity of our evaluation in small vessel detection. The non-vascular landmarks were placed densely in the vicinity of the vessel’s boundary and CSF region where over segmentation and false positive detection are most likely to occur. Representative examples of vessel and background landmarks from a young and aged control subjects are shown in (**Figure 2** (1) A-1, A-2), and (**Figure 2** (1) B-1, B-2), respectively.

**Figure 2.**
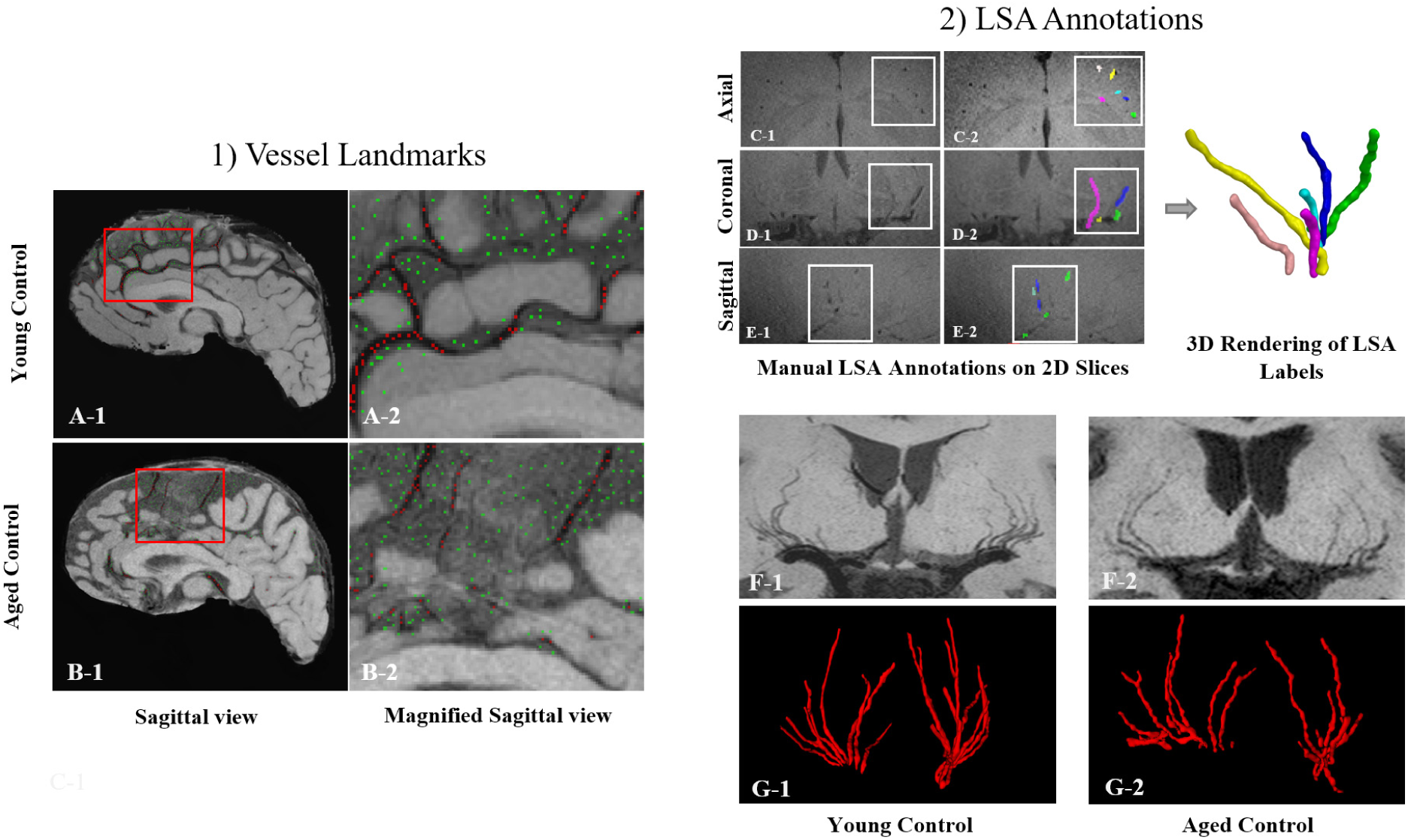
Dataset for method validation. (1) Vessel and background landmark annotation on a selected sagittal slice of two representative subjects, where (A-1) is a healthy 29-year-old male, and (B-1) is a healthy 64-year-old female. Magnified view of vessel landmarks (red) and background points (green) of the two subjects are shown in (A-2) and (B-2), respectively. (2) Manual lenticulostriate artery (LSA) annotation by experts using ITK-SNAP. For an axial (C-1), coronal (D-1), and sagittal (E-1) slice of a Black-blood MRI, manual delineated labels are on the axial (C-2), coronal (D-2) and sagittal (E-2) view are plotted. Additionally, minimum intensity projections of high-resolution 3T MRI of LSAs for a healthy 25-year-old male (F-1), and a healthy 67-year-old female (F-2) are shown together with their matching 3D rendering of LSA labels in (G-1) and (G-2), respectively.

To evaluate the segmentation accuracy of the candidate methods, the vessel segmentation was obtained by thresholding the vesselness map generated by each method, where the threshold value was carefully estimated by the Receiver Operating Characteristic (ROC) analysis using vessel landmarks of the validation dataset. Afterwards, the candidate methods with optimized parameters were tested on the independent LSA-30 dataset, where the LSAs were delineated manually by an experienced neuroradiologist. For an objective assessment of the segmentation performance, the following metrics were employed:

- Sensitivity (SEN) = TP/(TP + FN).
- Specificity (SPE) = TN/(TN + FP).
- Accuracy (ACC) = (TP + TN)/(TP + TN + FP + FN).
- Average Hausdorff distance (AVD)

where TP is true positive, FP is false positive, TN is true negative, and FN is false negative. The AVD metric is a widely used performance measure for calculating the distance between two point sets. In medical image segmentation, AVD is utilized to compare the distance between ground-truth and segmentation results, hence, smaller values indicate better performances (Taha and Hanbury, 2015). In this study, the AVD metric was calculated only for LSA segmentation results since we have complete annotation of these vessels.

### 2.4 Vessel Classification using Anatomical Masking

The vessels of interest in this study were small artery, arterioles, venules and small veins which were detected by Hessian-based methods applied on skull-stripped high-resolution black-blood MRI (0.5mm isotropic). The optimized scale (vessel radii) range for the study cohort was empirically estimated as [0.1-0.4] voxel (i.e., 50-200um) based on ROC analysis as described in section 2.3. To obtain the small vessels’ mask, the resultant vascular structure (or BVM) of skull-stripped high-resolution black-blood MRI was classified. More precisely, apart from a complete small cerebrovascular network, the final BVM included full or partial detection of superficial veins, dural sinuses (include superior sagittal sinus, straight sinus, transverse sinus, etc.) and middle cerebral artery (MCA) (**Figure 3** (a)). To separate these vessels, a region of interest (ROI) was defined via anatomical regions of MPRAGE images using the Freesurfer ASEG map. As shown in **Figure 3**, dural sinuses form the terminal of superficial venous system and distribute without closely accompanying arteries (Rhoton, 2002). Therefore, to isolate dural sinuses, superficial arteries and veins, a tentative mask (V_ten_) was generated by morphological erosion of the brain mask (consist of WM, GM, and CSF). Afterwards, to exclude regions with larger arteries and highly variable small vessel detection due to low SNR or imaging artifacts, the final ROI (**Figure 1**(e)) was constructed by eliminating cerebellum, brainstem, optic-chiasm and ventral regions from V_ten_. The ROI mask was then transformed to the black-blood image space using the affine transformation to the MPRAGE image as will be described in **Section** 2.6.

**Figure 3.**
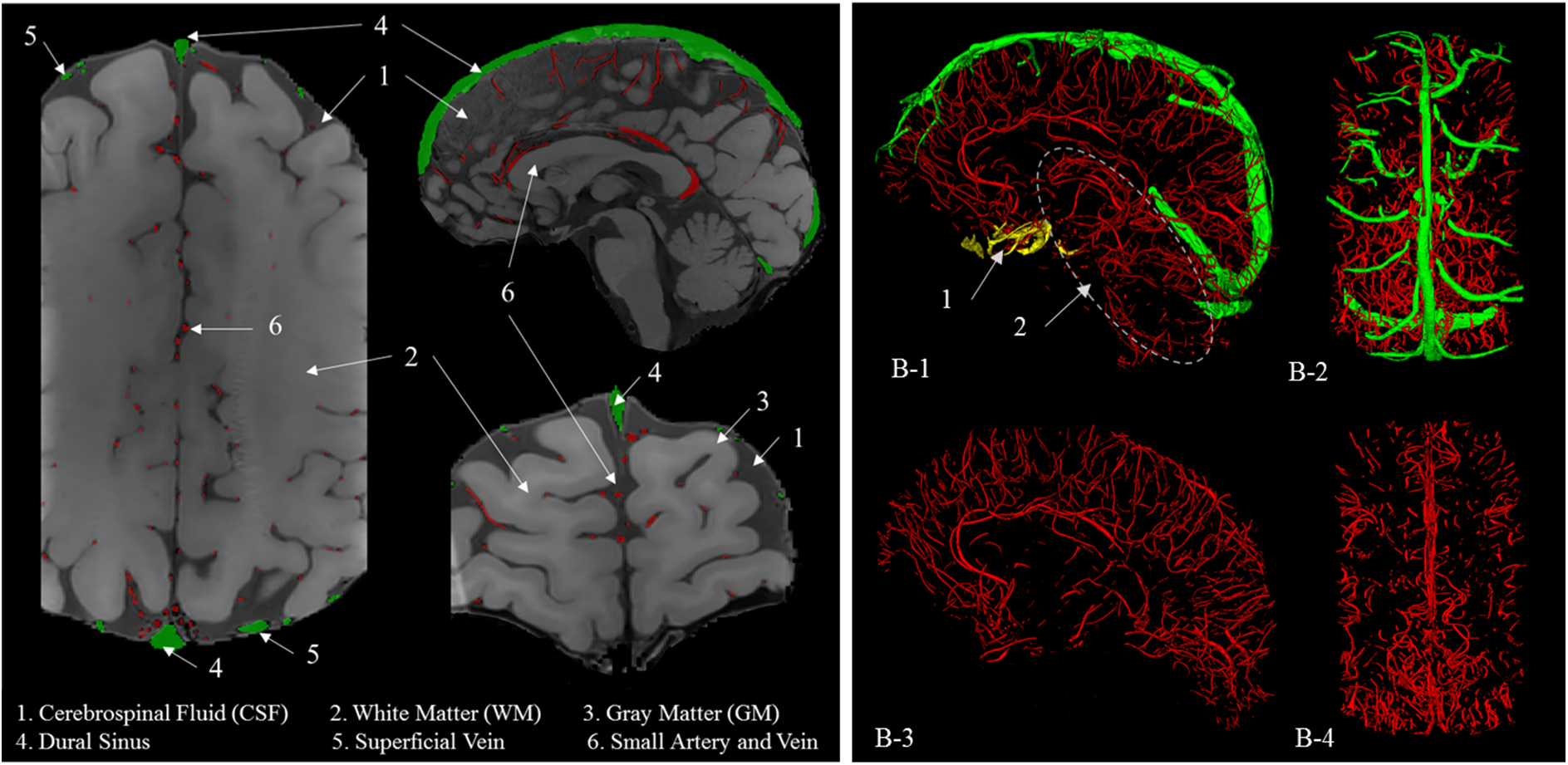
Spatial distribution of arteries, veins, and dural sinuses in high-resolution black-blood MRI. (a) Sagittal, coronal, and axial slices with overlayed vessel annotations. (b) 3D-rendering of vascular networks in (a); in (B-1, B-2) vessels are color coded to show vascular classification, where red-colored vessels are small arteries and veins, green-colored vessels are dural sinuses, and yellow-colored vessels are MCAs. To obtain the final masked small vessels (B-3, B-4), dural sinuses and brainstem and cerebellum regions ((1) and (2) highlighted in B-1) were excluded.

### 2.5 Vessel Density Calculation

Vessel density images (VDI) were calculated to allow localized comparison of vascular changes across individuals. This step is essential since the spreading pattern of small vessels in high resolution black-blood MRI varies among different subjects, which makes the direct comparison of cerebral small vessels at voxel-level a difficult task. To overcome this challenge, we computed VDI of each subject by diffusing the content of BVM to the entire image volume. More precisely, VDI was a smooth vessel map obtained by a convolution of the BVM with a 3D kernel. The window size of the kernel was empirically set as [30 30 30] voxels for all subjects.

Representative VDI examples of a young and aged control subject are shown in **Figure 4**. In the VDI representation, red regions denote high vessel density associated with dense vessel distribution. Blue regions indicate lower vessel density. According to the VDI representation, we can observe sparser vessel distribution in the aged subject (**Figure 4**(b)) and a denser vessel network in the young subject especially in frontal regions (**Figure 4**(a)).

**Figure 4.**
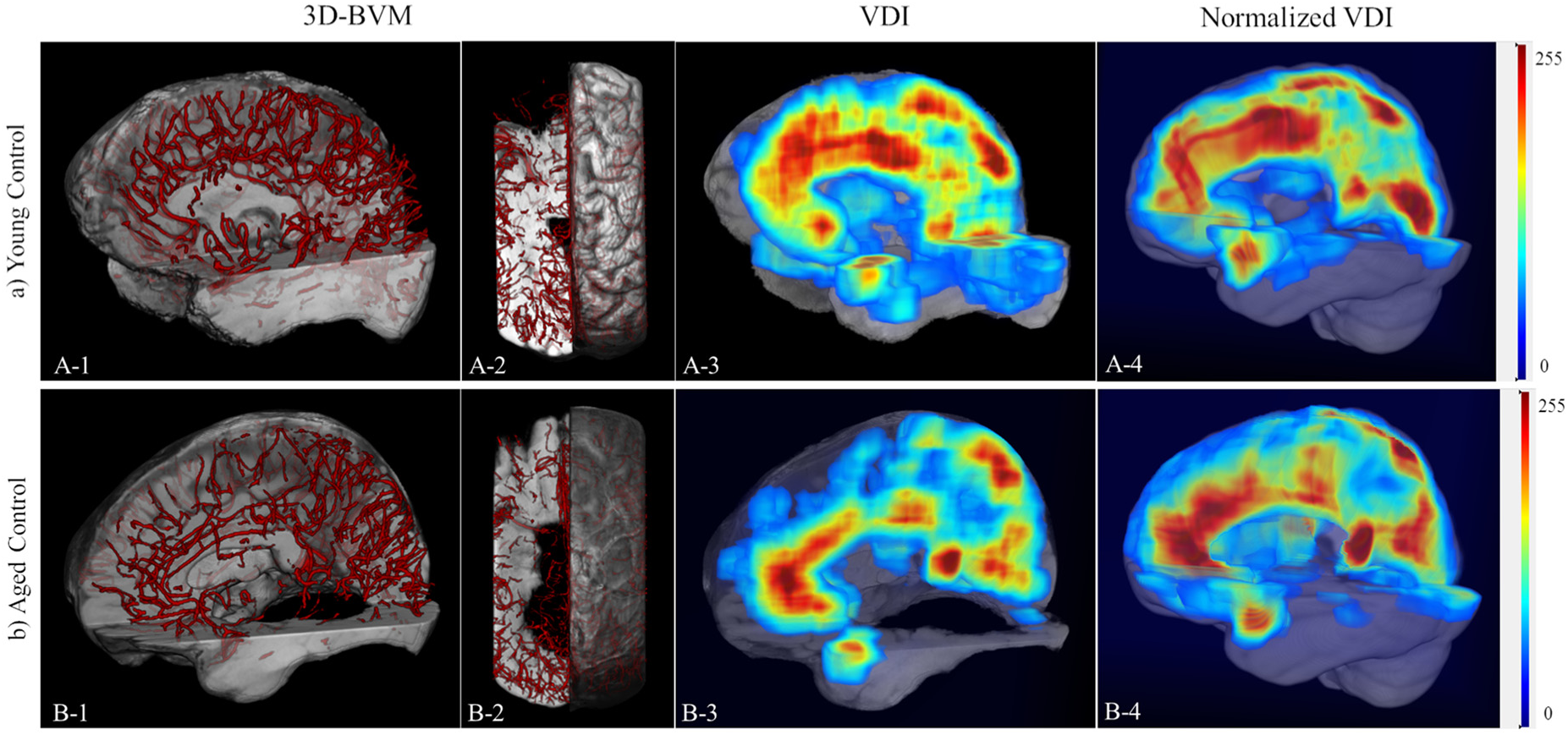
Representative examples of vessel density image (VDI) from two control subjects. (a) 3D-rendering of the BVM of a young control subject (A-1, A-2). (A-3) Corresponding VDI of the BVM in (A-1, A-2) before (A-3) and after normalization (A-4). (b) 3D-rendering of the BVM of an aged control subject (B-1, B-2). (B-3) Corresponding VDI of the BVM in (B-1, B-2) before (B-3) and after normalization (B-4).

### 2.6 Mapping Vessel Density Across Subjects

Due to anatomical variation, different brain size and MRIs’ orientations among subjects, it is required to normalize the subjects’ VDIs into common space to enable cross-subject comparison at voxel-level. To this end, we proposed a two-step registration method (supplement **Figure S2**). In the first step, to robustly co-register skull-stripped black-blood MRI and MPRAGE image pairs, a 3D affine registration with 12 landmark points was performed (supplement **Figures S2**(a), and (b)) using the Elastix software (Klein et al., 2010). The landmarks were manually selected from the cerebrum, cerebellum, and brainstem regions to guide the co-registration between the two 3D volumes with different field of views, where the black-blood MRI has near-whole brain coverage while the MPRAGE scan has full brain coverage. In the second step, we computed the nonlinear warp between the MNI152 Atlas (common space in this study) and the MPRAGE image (supplement **Figures S2** (c), and (e)). By combining the affine and nonlinear warp, we obtained the final transformation from the black-blood MRI to the MNI152 space.

To pool the VDIs from all subjects into the MNI152 space for statistical analysis, VDI was first reversely transformed to MPRAGE space using previously computed affine transformation (**Figure S2** (Step-1)), and then non-linearly transformed to the MNI Atlas using a B-spline transformation (**Figure S2** (Step-2)). After non-linear registration, all VDIs were normalized and transformed into the common space. Then, two-sample t-test was computed to compare VDIs of the two groups at voxel-level and statistical threshold was set to p<0.05 two-tailed (uncorrected).

### 2.7 Regression Analysis between Vessel Density and Cognitive Scores

A prominent behavioral phenotype of cSVD is early executive dysfunction manifested by impaired capacity to use complex information, to formulate strategies, and to exercise self-control, with less pronounced episodic memory deficits compared to AD patients (Wallin et al., 2018). Therefore, we used a composite score of executive function (EF) developed using IRT (Staffaroni et al., 2020a) which offers advantages such as better reliability, fewer statistical comparisons, and improved power to detect longitudinal change with smaller sample sizes (Gibbons et al., 2012; Staffaroni et al., 2020b). IRT EF composite scores were calculated for the 12 aged subjects (age range = [62, 81], Age mean±SD (years) = 68.2±7) who underwent neuropsychological assessment. A correlation analysis was performed at each voxel between vessel density image and neurocognitive scores including IRT EF composite score, IRT-Z EF score normalized by age, gender and education, and MoCA score which reflects gross cognition. The resultant correlation map was thresholded at p<0.05 (two-tailed, uncorrected) to detect the voxels with a significant association between vessel density and cognition scores.

## 3. Results

### 3.1 Validation Results of in vivo MRI

#### 3.1.1 Scale and threshold parameters estimation

The scale range of the candidate hessian-based methods was estimated as [0.1-0.4] with a step-size of 0.1 based on the vascular anatomy within ROI of the study cohort and was verified by the validation cohort with vessel landmarks. The optimal threshold for each filter (e.g., Frangi, Sato, Jerman) was determined by a point on the ROC curve closet to the optimal classifier (i.e., upper left corner of ROC plot). To robustly estimate this threshold, we used bootstrapped samples of the validation cohort with 1000 iterations and performed ROC analysis, where the scale range was fixed to the estimated value range of [0.1-0.4] voxels and the search window for the threshold value was defined as [0.001-0.2] with the step size of 0.001.

#### 3.1.2 Performance evaluation using vessel landmarks and LSAs

Evaluation and comparison of the multiscale filters with three enhancement functions: Frangi (Frangi et al., 1998), Sato (Sato et al., 2000) and Jerman (Jerman et al., 2016) were performed based on the annotated vessel landmarks of high-resolution black-blood MRIs (**Figure 5**) and tested using manual segmentation of LSAs (**Figure 6**), respectively. For all three methods, the scale range of multiscale Hessian-based methods was set to the target range of [0.1-0.4]. Utilizing the ROC analysis, the optimal threshold value was identified as 0.072, 0.005 and 0.15 for Jerman, Frangi and Sato’s filters, respectively. The three different thresholds for the candidate methods were due to the difference in their enhancement functions that results in vesselness maps with various range of values. We expect the winning method to generate more robust and generalizable vessel segmentation for all small vessels including LSAs with the same parameter setting.

**Figure 5.**
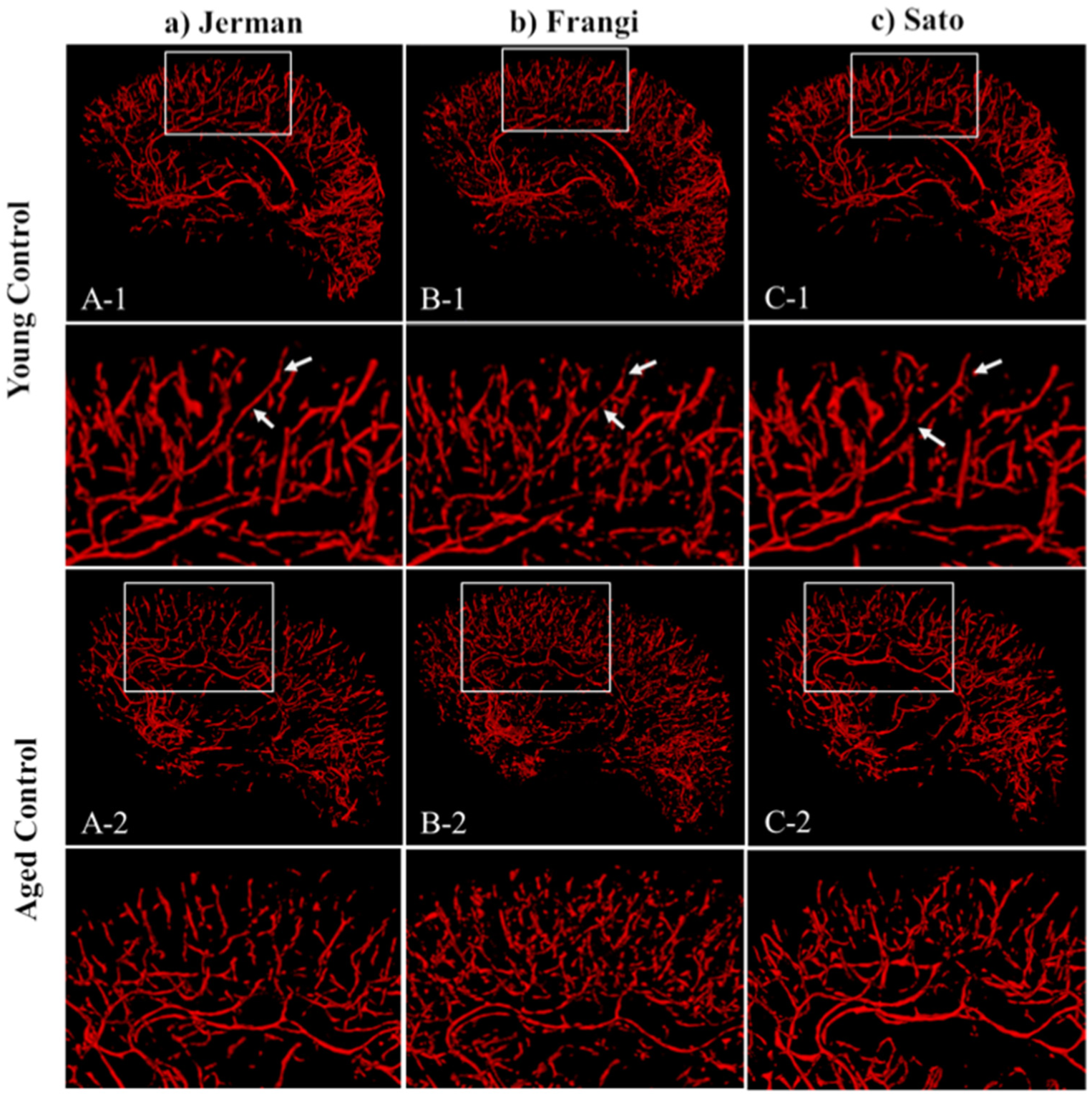
Performance comparison of Hessian-based methods for small cerebral vessel segmentation. (Row-1) 3D-rendering result of a young control subject (Female, 25 years old) by (a) Jerman, (b) Frangi, (c) Sato filter with optimized parameters. (Row-2) Magnified view of the annotated white box in Row-1. (Row-3) 3D-rendering result of an aged control subject (Female, 70 years old) by (a) Jerman, (b) Frangi, (c) Sato filter with optimized parameters. (Row-4) Magnified view of the annotated white box in Row-2. Overall, Jerman filter has more uniform and continuous vessel segmentation result for both young and aged subjects.

**Figure 6.**
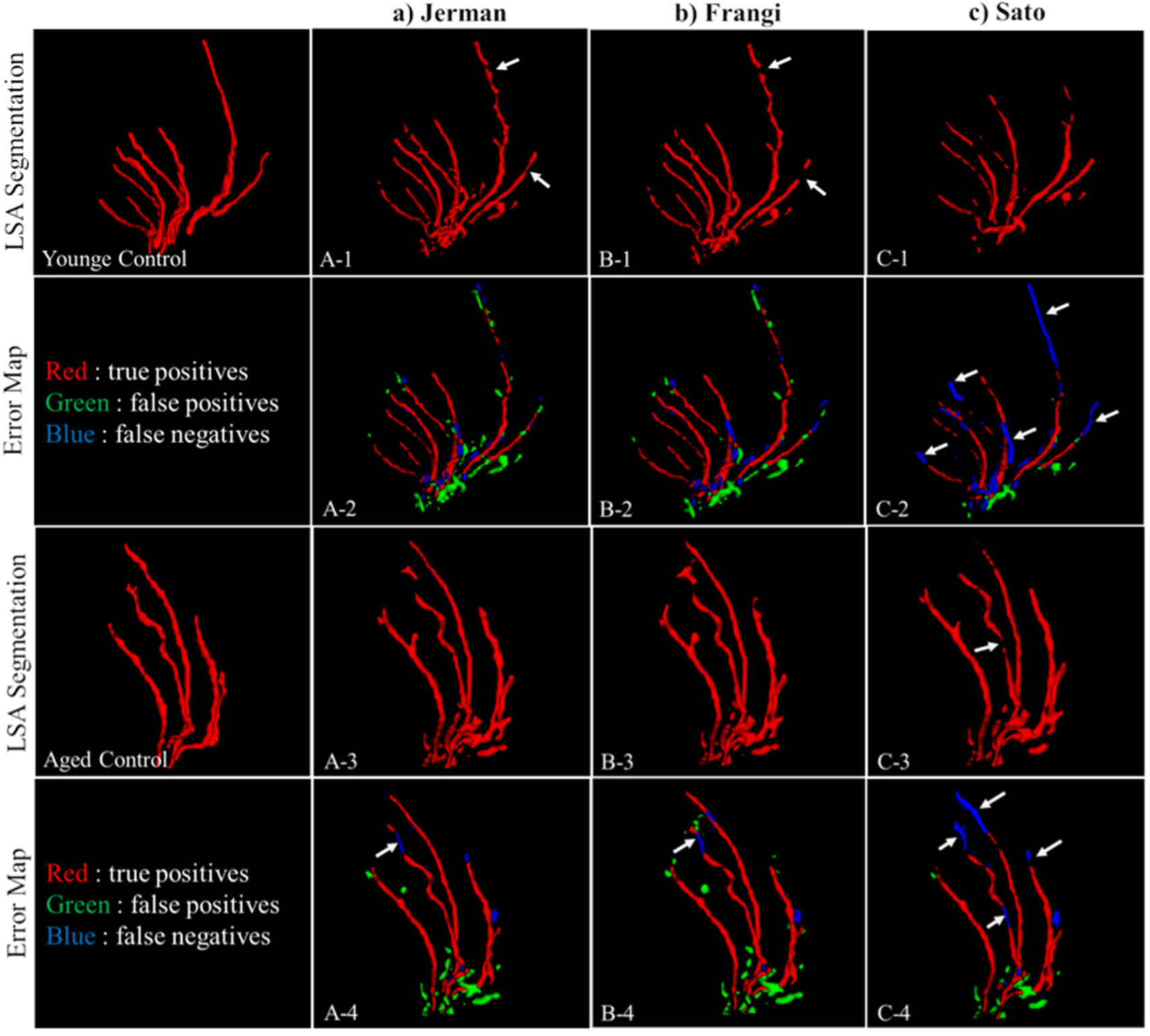
Performance comparison of Hessian-based methods for LSA segmentation of the same subjects in Figure 5. 3D-rendering of the LSA manual segmentation (column-1), and the segmentation result of the young (Row-1) and aged (Row-3) subject by (a) Jerman, (b) Frangi, (c) Sato filters with the same parameters as in Figure 5. (Row-2, Row-4) Corresponding error map of the segmentation results in (Row-1) and (Row-3), respectively.

Qualitative comparison of ROI vessels using the candidate methods of a young and aged subject is demonstrated in **Figure 5**. The results show that Jerman filter (**Figure 5**(a)) generated more robust, uniform and continuous vessel segmentation results compared to the other methods. Moreover, Frangi results had smaller, disconnected components in the vascular network segmentation of both young (**Figure 5**(b)(B-1) and aged (**Figure 5**(b)(B-2)) subjects. With Sato’s method, the resultant BVM had more discontinuity compared to Jerman’s and Frangi’s methods. For further assessment of the candidate methods, the output of LSA’s segmentation of the same young and aged subjects are demonstrated in **Figure 6**. The ground-truth of LSA delineation for both subjects are illustrated in **Figure 6**(Column-1). Jerman’s (**Figure 6**(a)) and Frangi’s (**Figure 6**(b)) methods had comparable results for LSA segmentation. The error map visualization of both methods shows that most of the vessels were detected correctly for the representative subjects (**Figure 6**(A-2, B-2, A-4, B-4). Jerman’s filter demonstrated less discontinuity compared to Frangi’s method in the young subject (white arrows in **Figure 6**(A-1, B-1)). Sato’s filter (**Figure 6**(c)) demonstrated lower detection rate compared to the other two methods mainly at the distal portion of the LSAs (white arrows in **Figure 6**(C-2, C-4)).

Quantitative results of the three methods based on the vessel landmarks of the validation dataset and LSA annotation of LSA-30 dataset are provided in **Table 1** and **Table 2**, respectively. According to **Table 1**, Jerman’s method showed the best performance in all three metrics including the AUC of ROC curves, sensitivity (SE) and specificity (SP). Table 2 of LSA validation results also shows that Jerman’s method outperformed the other two methods based on the four metrics including AUC, SE, SP and AVD.

**Table 1.** Summary of evaluation metrics for small vessel segmentation using vessel landmarks of validation dataset equally sampled from young control, aged control, and aged subjects with vascular risk factors.

| Method | ACCU | SE | SP |
| --- | --- | --- | --- |
| Jerman | 0.999 | **0.873** | 0.998 |
| Frangi | 0.994 | 0.844 | 0.994 |
| Sato | 0.995 | 0.749 | 0.995 |

**Table 2.** Summary of evaluation metrics for LSA segmentation using manual annotation of young and aged subjects.

| Method | AUC | SE | SP | AVD |
| --- | --- | --- | --- | --- |
| Jerman | **0.83** | **0.66** | 0.99 | **8.52** |
| Frangi | 0.81 | 0.63 | 0.99 | 10.58 |
| Sato | 0.73 | 0.46 | 0.99 | 18.64 |

### 3.2 Qualitative Results in Young and Aged Subjects

Representative examples of 3D small vessel segmentation of high-resolution black-blood MRI using Jerman filter with optimized parameters are demonstrated in **Figure 7**. Four cases were selected from the young control (**Figure 7**(a)) and aged control (**Figure 7**(b)) group respectively. Overall, by qualitative comparison we can see that young control subjects had significantly higher vessel density compared to the aged group.

**Figure 7.**
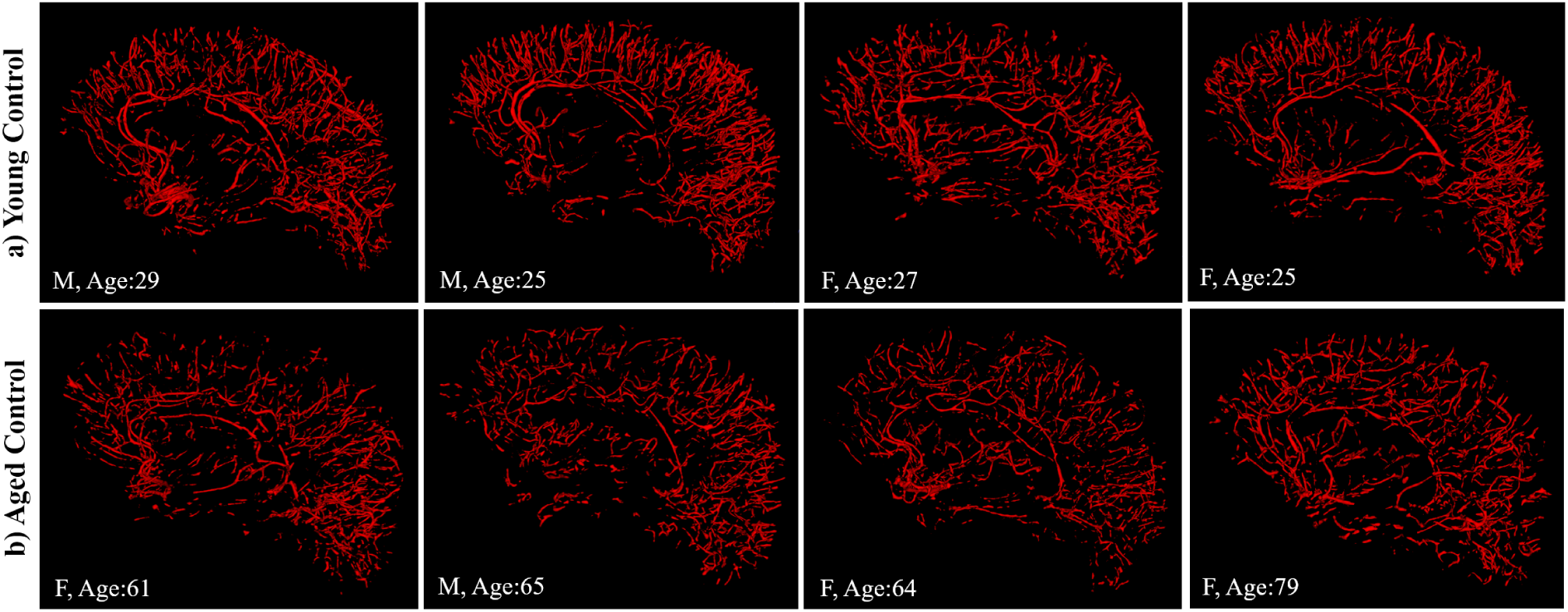
3D visualization of small vessels segmentation using Jerman method with optimized parameters. (a) young control, (b) aged control groups (second row) have sparser vessel network compared to young group (first row).

### 3.3 Quantitative Results

#### 3.3.1 Global Mean Vessel Density

**Figure 8** illustrates the comparison of mean small vessel density between the YC and AC groups. To account for variation of brain size across different individuals, small vessels’ volume was normalized by the ROI size. More precisely, normalized vessel density measure of each subject equals to (small vessels volume/ROI volume), where ROI is defined in **Section** 2.4. As shown in **Figure 8**, the result of two-way student t-test demonstrated that the YC group had significantly higher vessel density compared to the aged group (p<0.05).

**Figure 8.**
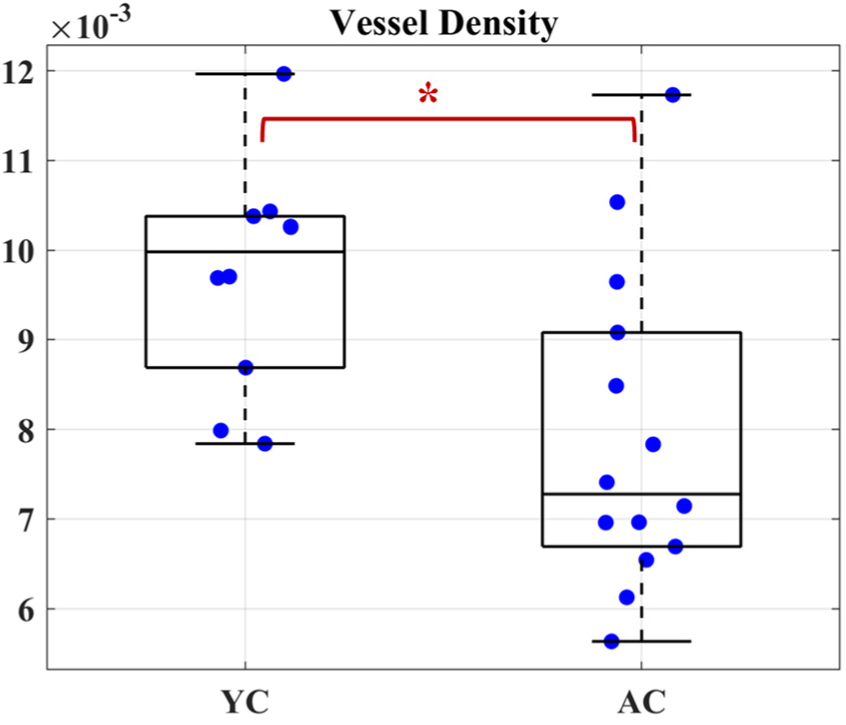
Comparison of mean vessel density of small cerebral vessels in young (age 22-33 years) and aged (age > 60 years) groups. * indicates significance p<0.05.

#### 3.3.2 Localized Vessel Density Analysis

##### 3.3.2.1 Group-wise Analysis of Age-dependent Vessel Density Changes in Normal Brains

**Figure 9** demonstrates the quantitative results of voxelwise comparison between the young and aged groups in the near-whole brain volume, where we projected the t-value maps onto a cortical surface in the MNI152 space for better visualization. We observed widespread significant reductions in vessel density in the aged group as compared to the young group. These regions mainly include medial, superior and middle frontal gyrus, precentral gyrus, precuneus and cuneus.

**Figure 9.**
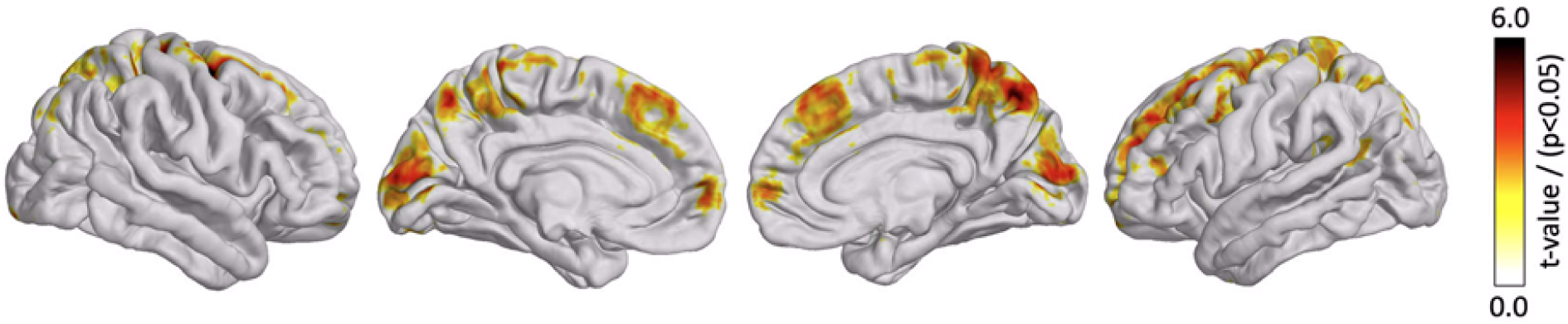
Areas with significant (p<0.05, uncorr.) higher vessel density in young controls (YC (N=10)) as compared to aged controls (AC (N=18)).

##### 3.3.2.2 Regression Analysis of Vessel Density Changes and Cognitive Scores

The normalized VDIs of twelve aged control subjects (age range = [62, 81], mean±SD (yrs) = 68.2±7) who underwent neuropsychological assessment were correlated with their cognitive and executive function scores (such as IRT EF, IRT-ZScore of EF, and MoCA) and the resultant correlation maps are shown in **Figure 10**. Consistent Associations across all cognitive scores were observed in posterior cingulate cortex (PCC), dorsolateral prefrontal cortex (dlPFC) as well as in medial temporal lobe.

**Figure 10.**
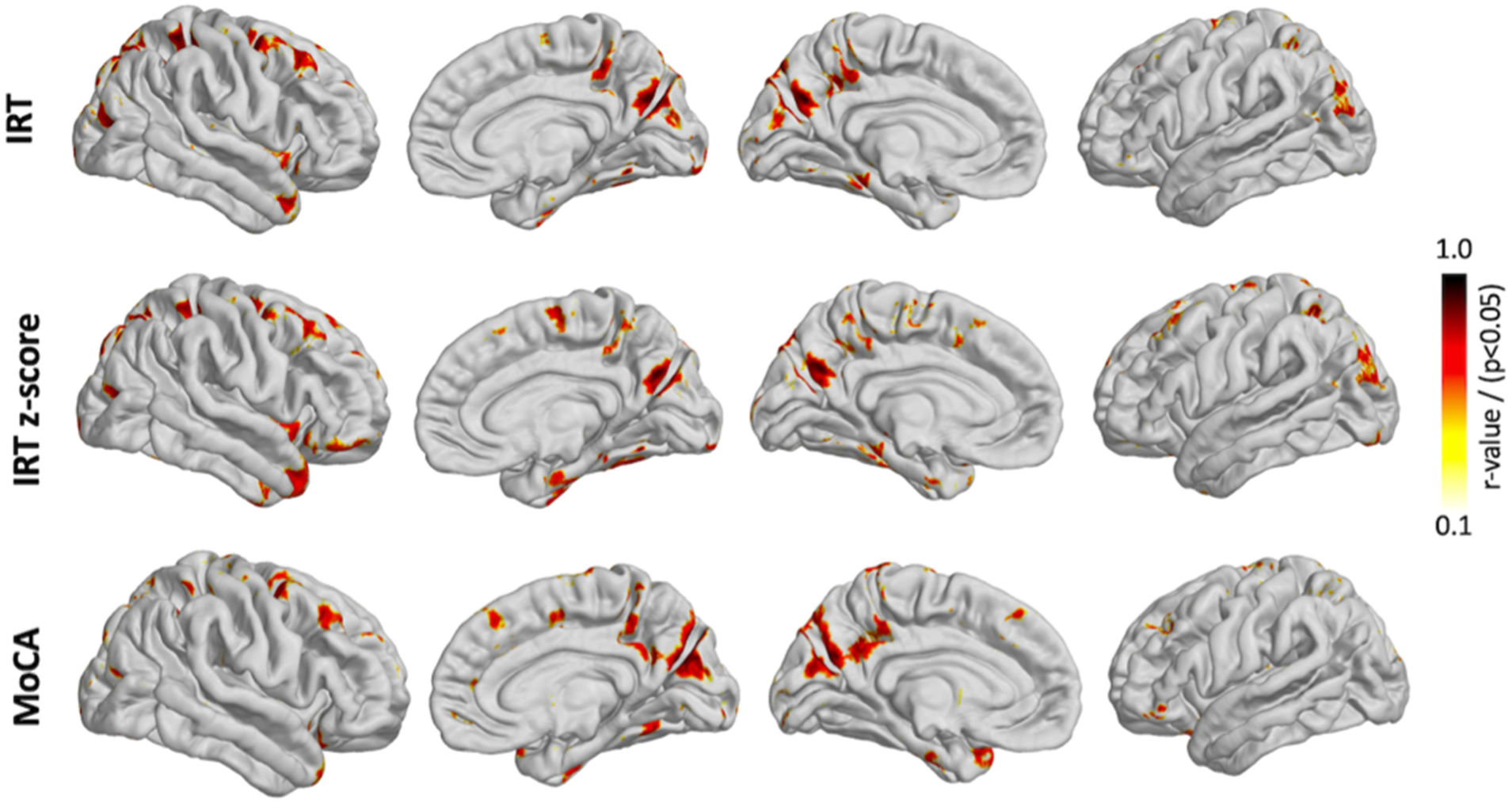
Localization of significantly associations between VDI of AC (N=12) subjects and their corresponding IRT score, IRT-ZScore score and MoCA score.

## 4. Discussion

In this study, we developed, optimized and validated a comprehensive 3D analysis framework for localized mapping of small vessel based on high resolution (isotropic ∼0.5mm) black-blood MRI with near whole-brain coverage of young and aged subjects at standard clinical magnetic field strength of 3T. In our experimental results, we evaluated the performance of the proposed framework in studying the small vessel density changes in normal aging, and in correlation with cognitive scores. Overall, we observed a decreasing pattern in the small vessel density by aging, and a positive correlation between small vessels density and cognitive scores such as IRT EF and MoCA scores.

### 4.1 Clinical Value of Whole Brain Small Vessel Density Mapping

To date, limited in vivo techniques are available to directly assess cerebral small vessels across the whole brain. Digital subtraction angiography (DSA), x-ray computed tomography angiography, and time-of-flight MR angiography have been applied to observe large cerebral vessels in clinical populations (Gotoh et al., 2012; Kammerer et al., 2017; Wardlaw et al., 2013b), but the sensitivity to small vessels such as the perforating and pial arteries is moderate at best and difficult to characterize and quantify at 3T. With the proposed 3D multimodal framework for analyzing high-resolution black-blood MRI, the automated segmentation enables both visualization and quantitative assessment of cerebral small vessels.

### 4.2 Segmentation and Quantification

This study was inspired by our previous work (Sarabi et al., 2020), where we developed a 3D retinal vessel density mapping approach for optical coherence tomography angiography (OCTA) and demonstrated its efficacy in localized detection of microvascular changes and monitoring retinal disease progression in different clinical applications such as normal aging and diabetic retinopathy (DR). Despite some similarities between the two studies, with respect to two-module framework design for fine-scale vessel segmentation and non-linear registration, there are still some fundamental differences in the method development. 3D cerebral vascular mapping in high resolution black-blood MRI images is more challenging compared to retinal vessels mapping due to relatively poor vessel contrast mainly in cerebrospinal fluid (CSF) regions, varying degrees of noise, inhomogeneous backgrounds, anatomical variations, and partial view of the brain in this study. To tackle these challenges, initially high-resolution black-blood MRIs underwent several preprocessing steps such as skull-stripping, bias correction and NLM denoising. NLM was selected for denoising since previous works (Chen et al. 2011, Zhang et al. 2014) demonstrated that this process preserves anatomical details while suppressing the noise. Since the microvasculature involved in cSVD includes small arteries/arterioles (∼10µm-1mm), capillaries (<10µm), and venules (∼10-50µm) (Charidimou et al., 2016), we employed multi-scale Hessian-based method for the purpose of vessel segmentation. More precisely, multiple popular Hessian-based methods (Frangi et al., 1998, Sato et al., 2000, Jerman et al., 2016) have been evaluated for the enhancement of small vessels using clinical data. Our evaluation results demonstrated that the Jerman filter was more robust to noise compared to other candidate methods, had higher and more uniform responses for small vessels not only at vessel center but also at vessel periphery. To robustly detect the small vessels, validation experiments were designed to determine the optimal scale and threshold value via ROC analysis using a validation dataset with annotated vessel landmarks. For localized small vessel density analysis across populations, first, a vessel density image (VDI) was calculated by diffusing the content of the vessel mask to the entire image volume. After that, we implemented a non-linear registration framework to pool the VDIs from all subjects into the common space (MNI-152 Atlas). To this end, initially skull-stripped black-blood MRI and MPRAGE image pairs were robustly co-registered using a 3D-Affine registration with 12 landmark points. Subsequently, a non-linear registration was performed between MPRAGE of each subject and MNI-152 Atlas using B-Spline method. By utilizing these two deformation fields, all VDIs were normalized to MNI-152 Atlas. Thus, together with the careful preprocessing of the image data, the localized mapping of vessel density in different subjects and patient groups enabled the analysis of region-specific changes across populations.

### 4.3 Aging and Association with Neurocognitive Scores

Our approach of vessel density mapping revealed decreasing small vessel density with aging which is consistent with literature (Sadoun and Reed, 2003; Watanabe et al., 2020). Age-related cerebral microvascular changes have been visualized using ultrasound-based microscopy in animals (Lowerison et al., 2022), while retinal vessel density can be quantified using OCTA in humans which was correlated with cognitive scores such as MoCA (Ashimatey et al., 2020; Sarabi et al., 2020). Laser scanning confocal 3D immunofluorescent microscopy of young and aging vascular beds in multiple mice and human organs, including brain tissue, has also demonstrated an age-dependent decline in vessel density (Chen et al., 2021). To the best of our knowledge, this is the first in vivo study on aging effects on cerebral small vessel density using MRI. We also explored associations between localized vessel density and neurocognitive scores including IRT EF and MoCA. IRT EF scores were chosen since early executive dysfunction has been shown to be a prominent behavioral phenotype of cSVD (Wallin et al., 2018). The IRT-based score used in the present study is sensitive to impairments in mild cognitive impairment (MCI), Alzheimer’s disease (AD), and frontotemporal dementia (Staffaroni et al., 2020b) and is associated with cerebral blood flow of middle cerebral artery perforator territory (Jann et al., 2021). In the present study, we found significant positive associations between regional vessel density and IRT EF scores in areas associated with executive functions such as dlPFC and PCC. In addition, we found a similar pattern of brain areas with significant positive associations between regional vessel density and MoCA scores (Nissim et al., 2016; Yokosawa et al., 2020). These promising data suggest that the proposed analysis framework may be useful as an early screening tool for cSVD to gauge overall health of cerebral small vessels. Furthermore, it could serve as a biomarker to monitor disease progression and/or response to interventions for vascular dementia.

### 4.4 Limitations of the Study

In a few subjects, near-whole brain coverage instead of full FOV caused wraparound of ear tissue signal into the image at the brainstem. Saturation bands should be placed more carefully to saturate the signal of tissue outside the FOV. Such high-resolution imaging is susceptible to motion artifacts that cause blurring or ghosting in the images. Motion was minimized by padding the subjects in the head coil and applying paper tape across the forehead skin for tactile feedback. Motion compensated 3D TSE-VFA sequences may provide a robust method for small vessel visualization and quantification (Hu et al., 2021). As a proof-of-concept study, we only included a small cohort of young and aged subjects, therefore no correction of multiple comparisons was performed for statistical analysis.

One potential confounding factor for segmenting small vessels on black-blood MRI is perivascular space (PVS) which appears gray on T1w TSE-VFA images. We have compared LSA and PVS in T1w and T2w TSE-VFA respectively, as shown in Figure S7 of (Ma et al., 2019), and the locations of LSAs did not match those of PVS in the basal ganglia area. In contrast to PVS that is primarily observed in white matter, the majority of segmented superficial small vessels are in gray matter, as shown in **Figure 3**. The relationship between small vessels and PVS in terms of their spatial distribution and quantity need to be evaluated in future studies.

## 5. Conclusions

We presented and evaluated a novel framework for segmenting and mapping brain small vessels from high-resolution black-blood images acquired at 3T. Using filter-based segmentations and non-linear registration, 3D mapping, and quantification of brain small vessels was demonstrated to be feasible. This framework may serve as a promising tool for localized detection of vessel density changes in patients with neurovascular diseases and in understanding the correlation between vessel density and cognitive scores.

## ACKNOWLEDGEMENT

This work was supported by National Institute of Health (NIH) grants UF1-NS100614, R01-NS114382, R01-EB032169, R01-EB028297, R01-EB022744, RF1-AG064584, P30AG066530, and P41-EB015922.

## Supplementary material

**Figure S1.**
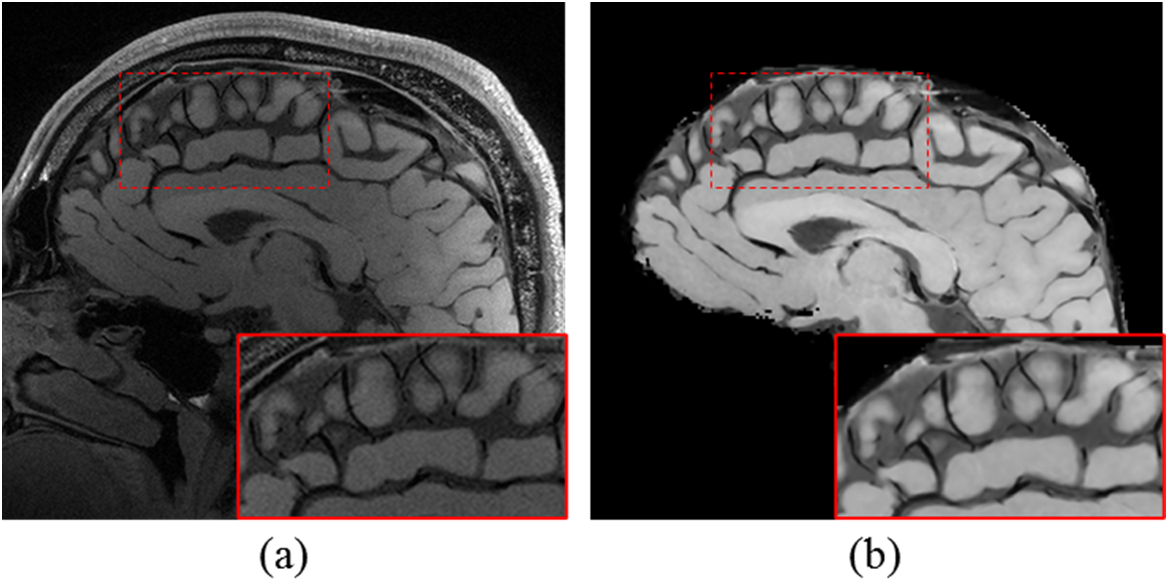
Example high-resolution black blood MRI preprocessing shown on a selected sagittal scan. (a) Raw image of a young control subject. (b) preprocessed image using skull-stripping, bias correction and NLM denoising.

**Figure S2.**
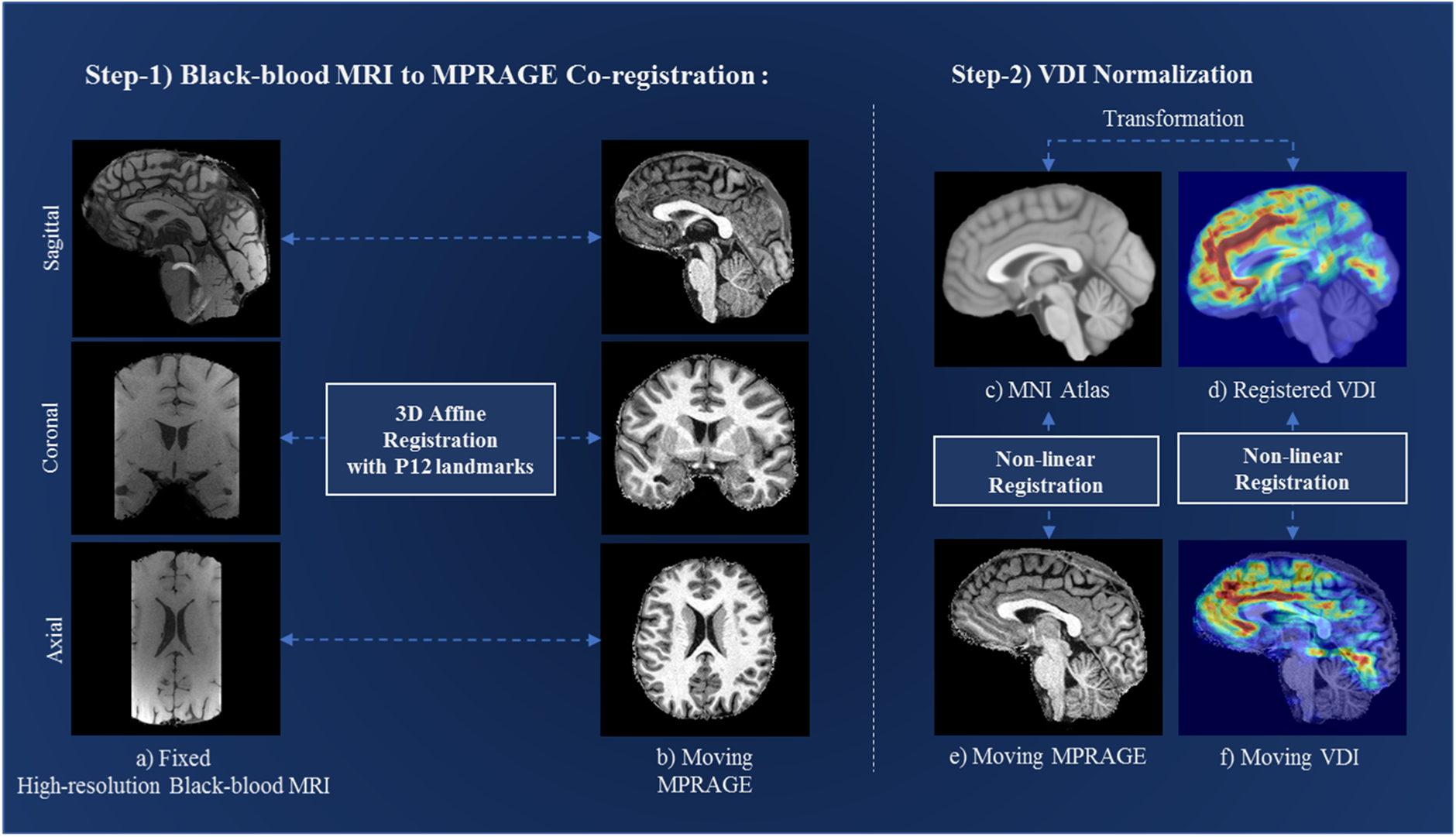
Flow chart of co-registration and VDI normalization. Step1: Co-registration of high-resolution black-blood MRI (a) and MPRAGE (b) image pairs, using 3D-Affine registration with 12 landmark points, where (a) is the fixed and (b) is the moving image. Step2) Nonlinear warp between the MNI-152 Atlas (c) and MPRAGE (e) image. VDI was first reversely transformed to MPRAGE space using affine transformation (f), and then non-linearly transformed to the MNI Atlas using a B-spline transformation (d).

## Notes

### Competing Interest Statement

The authors have declared no competing interest.

### Summary of Updates

A new funding source P30AG066530 has been added to the acknowledgement section.

